# A synthetic peptide library for benchmarking crosslinking mass spectrometry search engines

**DOI:** 10.1101/821447

**Authors:** Rebecca Beveridge, Johannes Stadlmann, Josef M. Penninger, Karl Mechtler

## Abstract

We have created synthetic peptide libraries to benchmark crosslinking mass spectrometry search engines for different types of crosslinker. The unique benefit of using a library is knowing which identified crosslinks are true and which are false. Here we have used mass spectrometry data generated from measurement of the peptide libraries to evaluate the most frequently applied search algorithms in crosslinking mass-spectrometry. When filtered to an estimated false discovery rate of 5%, false crosslink identification ranged from 5.2% to 11.3% for search engines with inbuilt validation strategies for error estimation. When different external validation strategies were applied to one single search output, false crosslink identification ranged from 2.4% to a surprising 32%, despite being filtered to an estimated 5% false discovery rate. Remarkably, the use of MS-cleavable crosslinkers did not reduce the false discovery rate compared to non-cleavable crosslinkers, results from which have far-reaching implications in structural biology. We anticipate that the datasets acquired during this research will further drive optimisation and development of search engines and novel data-interpretation technologies, thereby advancing our understanding of vital biological interactions.

## Introduction

Chemical crosslinking combined with mass spectrometry (XL-MS) is frequently used to gain structural information on proteins and protein complexes [1, 2]. In a crosslinking experiment, a reagent forms covalent bonds between specific amino acid side-chains that are in close spatial proximity, thus revealing distance restraints between residues, and hence interaction sites within a protein or between different proteins [3, 4]. Initial applications of these techniques were limited to small proteins and protein complexes. More recently, due to major methodological and technological developments, XL-MS has been applied to living cells to investigate protein interactions and topological structures at the proteome-wide level [5–8].

For studies on small proteins and protein complexes, standard crosslinking reagents disuccinimidyl suberate (DSS), bis-(sulfosuccinimidyl) suberate (BS3) or bis- (sulfosuccinimidyl) glutarate (BS2G) are typically used which react with lysine residues and N-termini of proteins, and to a lesser extent with serine, threonine and tyrosine. Several complications arise during the identification of crosslinked peptides with such reagents. Measurement of the intact mass of the crosslinked moiety, rather than the mass of the individual peptides, quadratically increases the number of possible peptide pairings to be searched with the number of peptides in the database (often referred to as the ‘n-squared’ problem), and is thought to reduce confidence in the assignment of crosslinked peptides [9]. With the aim of overcoming this limitation, MS-cleavable crosslinkers such as disuccinimidyl dibutyric urea (DSBU) and disuccinimidyl sulfoxide (DSSO) were introduced [10, 11]. This class of reagents contain MS-labile bonds at either side of a functional group within their spacer regions that can be selectively and preferentially fragmented prior to peptide backbone cleavage during collision induced dissociation (CID) or higher-collision induced dissociation (HCD). Gas-phase cleavage of the crosslinking reagent during tandem MS enables MS3 acquisition methods, which facilitate peptide sequencing using traditional database search engines due to circumvention of the n-squared problem. Additionally, cleavable cross-linkers generate diagnostic ion doublets during MS2, which are required for the use of novel database search engines such as XlinkX and MeroX, that provide means for proteome-wide XL-MS studies [12, 13].

Another critical challenge during XL-MS lies in estimating the error rate in a search for crosslinked peptides. In protein identification by mass spectrometry an FDR (false discovery rate) method is often used, whereby incorrect decoy sequences added to the search space correspond with incorrect search results which might otherwise be deemed correct. This allows the estimation of how many incorrect results are in a final data set. In XL-MS, the FDR estimation is complicated by the fact that every match is a combination of two peptides, each with its own probability to be false. Further, false discovery rates can be estimated at different points during data analysis, for example protein or residue pairs, and misuse of these approaches can lead to a higher error in the results than is targeted in the search [9].

Owing to the versatility of XL-MS techniques, many algorithms have been developed to identify crosslinks from mass spectrometry datasets. For example, algorithms used in conjunction with non-cleavable, non-labelled crosslinkers include (but are not limited to) pLink [14], StavroX [15], Xi [16], and Kojak [17]. For MS-cleavable reagents, XlinkX [12] and MeroX [13] are often used. pLink, StavroX, Xi, MeroX and XlinkX have an in-built FDR calculator, whereas Kojak relies on external tools to estimate FDRs such as Percolator [18] or PeptideProphet [19] which is incorporated into the trans proteomic pipeline [20].

Crosslinks are often validated by mapping them onto crystallographic models and measuring the distance between the α-carbon atoms of the crosslinked residues [10, 11, 21, 22]. Distances that are within a defined cut-off point are considered to be true, and those that exceed this distance limit, and are therefore in disagreement with the structural model, are regarded as false positives. One limitation of such methods to test bioinformatic approaches is that the formation of non-specific crosslinks during the experimental procedure cannot be fully eliminated, and such a crosslink would wrongly be defined as a false positive. Furthermore, such an approach often under-estimates FDR since the crosslinks that are evaluated are limited to those that are formed between peptides contained in the same protein model [23]. This lack of a generally accepted method on how to control for falsely identified crosslinks may have severe implications that are far-reaching in biological research.

Several elegant benchmarking approaches were utilised by Chen et al. [24] during the evaluation of pLink 2. Simulated datasets, synthetic datasets [25], ^15^N metabolically-labelled datasets and entrapment datasets [17, 26] were used to compare the precision and sensitivity of pLink 2 compared to other search engines. The synthetic dataset was obtained by taking 38 synthetic peptides derived from the sequence of the UTP-B protein and crosslinking them in one reaction. Data were searched against increasingly larger databases, and the sensitivity and precision of pLink 1, pLink 2 and Kojak was reported.

To date, comprehensive assessment of the accuracy and sensitivity of XL-search algorithms have been hindered by a lack of ‘ground truth’ data that can be used to determine if crosslinks were correctly or incorrectly assigned. To overcome this problem, we have constructed a synthetic library of crosslinked peptides, which for the first time allows the unambiguous discrimination of XL-MS spectra that are correctly, incorrectly, and not identified across all search engines developed for the identification of crosslinks from the MS data.

## Results

### Design of the crosslinked peptide library

The crosslinked peptide library was based on the amino acid sequences of tryptic peptides from *S. pyogenes* Cas9 and was chemically synthesised according to Figure 1. 95 peptides were selected for synthesis that were 5-20 residues long, with single missed cleavage that is a lysine residue for crosslinking purposes. To control for unintended side-reactions of primary amine groups in the process of peptide crosslinking, any C-terminal lysine residues were incorporated into the peptide as epsilon-azido-L-lysine, and the N-terminus of the peptide was protected from crosslinking with a biotin group covalently linked to a generic ‘linker’ peptide sequence YGGGGR, followed by the library peptide sequence (Figure 1a). Subsequently, the only crosslinker-reactive site in each peptide at the time of crosslinking was one single lysine residue. Peptides were crosslinked with the gradual addition of crosslinking reagent to favour the formation of crosslinks (as opposed to monolinks that would likely form if all the XL reagent was added at once). The crosslinked peptides were then treated with trypsin overnight to enzymatically cleave the N-terminal biotin-linker peptide region, and subsequently treated with TCEP to reduce the C-terminal lysine azido-group to an amine. After trypsin digestion, the most abundant species in the solution was the biotin-linker, which strongly interfered with MS measurements, limiting the amount of material that could be loaded onto the analytical column. The biotinylated peptide species was therefore removed with streptavidin beads (figure 1b), and the resulting mixture exclusively comprised of crosslinked and monolinked tryptic peptides. To create a challenging dataset for the assessment of XL search algorithms, the synthetic peptides were divided into 12 groups for crosslinking, which were then combined prior to analysis *via* LC-MS. This theoretically gives rise to 426 potential crosslinks. It can be inferred that crosslinks identified between two peptides of the same group are correct, whilst those identified between two peptides from separate groups are known to be false positives (Figure 1c).

**Figure 1;.**
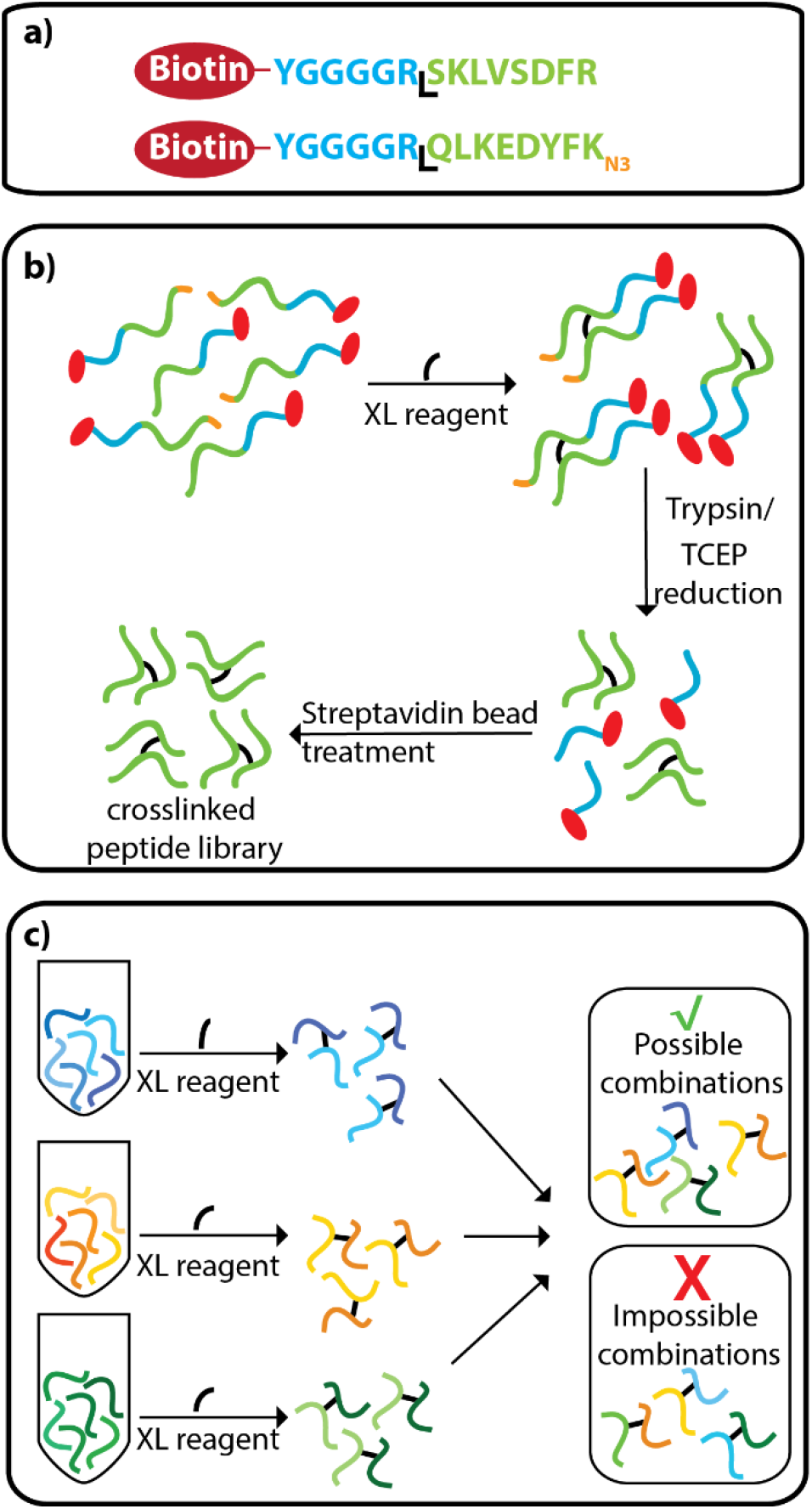
Design of the crosslinked peptide library. (a) the N-terminal amine group is protected from crosslinking with a biotin group that is followed by a linker region with a tryptic cleavage site. C-terminal lysine residues are incorporated with an azide group to prevent crosslinking to this site. (b) peptides are crosslinked, treated with trypsin, and the azide groups on the C-terminal lysine residues are reduced to an amine. Biotin-linker groups are removed from the solution with streptavidin beads. (c) the synthetic peptides are crosslinked in separate groups and combined prior to LC-MS analysis.

### Assessment of search engines for non-cleavable crosslinkers

The DSS-crosslinked peptide library was measured via LC-MS/MS and the resulting data were analysed with pLink, StavroX and Xi (Figure 2). Figure 2a shows the number of crosslink spectrum matches (CSMs) identified by the different search algorithms that correspond to correct (black) and incorrect (grey) crosslinks. Values are an average of three technical replicates, and the error bars represent the standard deviation. For all algorithms, the data were searched against the sequence of S. pyogenes Cas9 and 10 additional proteins (supplementary information) and the results were filtered to an estimated 5% FDR at the CSM level. pLink identifies the highest number of correct CSMs (678) and has a calculated FDR of 4.3%. StavroX and Xi identify a lower number of correct CSMs (410 and 512 respectively), with lower calculated FDR values of 2.4% and 2.5% respectively. The Venn diagrams display the overlap of scans from which CSMs are derived by the three algorithms, corresponding to correct (left) and incorrect (right) crosslinks. Out of 682 spectra that are matched to correct crosslinks by at least 1 search engine in the first technical repeat, 275 are matched by all 3, and almost all the spectra (639) are matched by pLink. Out of 45 scans that are matched with incorrect crosslinks by at least one algorithm, 27 are matched by pLink. One scan is matched to the same, incorrect crosslink by all 3 algorithms. Upon manual inspection of the spectrum, we found that the incorrect isotope peak is assigned to the mass of the precursor by the mass spectrometer.

**Figure 2;.**
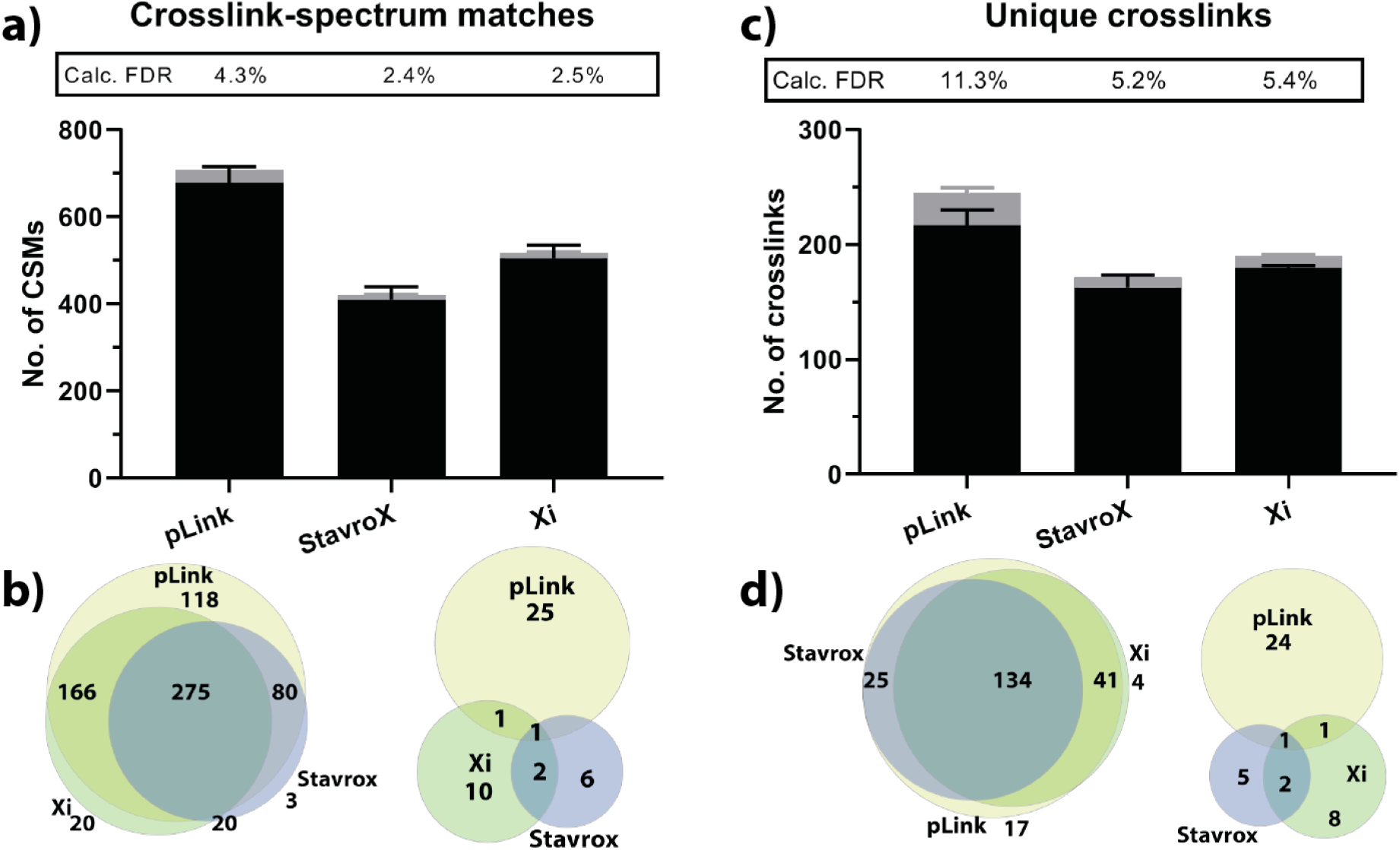
(a) number of CSMs that correspond to correct (black) and incorrect (grey) crosslinks identified by three search algorithms with incorporated FDR estimation. Results were filtered to an estimated 5% FDR, and the calculated FDR is given for each algorithm. Error bars correspond to the standard deviation between three technical replicates. (b) agreement of correct and incorrect CSMs between pLink, StavroX and Xi for one technical repeat. (c) number of correct unique crosslinks (black) and incorrect crosslinks (grey). (d) overlap of correct and incorrect crosslinks for one technical repeat. Values for figures a and c are given in Supplementary Tables 2 and 3.

The number of correct and incorrect unique crosslinks identified by the 3 algorithms are shown in Figure 2c. In the case of all algorithms, the calculated FDR of the unique crosslinks is higher than that of the CSMs. Whilst the FDR of the CSMs identified by pLink is 4.3%, this is propagated to 11.3% in the case of the unique crosslinks. This is due to CSM redundancy. For the correct crosslinks there is an average of 2.9 CSMs per crosslink. For the incorrect crosslinks, this ratio is 1.0. This effect is similar for StavroX and Xi that have CSM redundancies of 2.4 and 2.7, respectively. Out of 221 crosslinks that are identified by at least 1 search engine in the first technical repeat, 217 are identified by pLink, 179 are identified by Xi, and 159 are identified by StavroX (Figure 2d).

The number of CSMs and crosslinks identified when the results were filtered to an estimated FDR of 1% are shown in Supplementary Figure 1. Calculated FDR values of the unique crosslinks are 6.6%, 1.1% and 1.6% for pLink, StavroX and Xi, respectively, with pLink identifying 207 correct crosslinks, StavroX identifying 102, and Xi identifying 152.

### Effect of validation strategies on crosslink results

The effect of different validation methods on the sensitivity and accuracy of crosslink assignment is shown in Figure 3. In all cases, the results were filtered to an estimated FDR of 5%. We started by validating Kojak results with different tools, namely PeptideProphet [19] and Percolator [18] (Figure 3a), since Kojak has no built-in validation method. We observed that PeptideProphet is a very stringent validation method that has a calculated FDR of 2.4%, and that sensitivity is very low compared to other algorithms; only 123 correct crosslinks were identified. Percolator was used for validation by considering all CSMs identified by Kojak, or by considering only the highest-scoring CSM for each species (Unique CSMs). Both methods allow a much higher degree of sensitivity, but this comes with the price of much lower precision. When all CSMs are taken into account 224 correct crosslinks were identified, which is the highest of all identification approaches, but with a highly elevated FDR of 32%. When only the Unique CSMs were used, 221 correct crosslinks were identified with a slightly lower FDR of 23%.

**Figure 3;.**
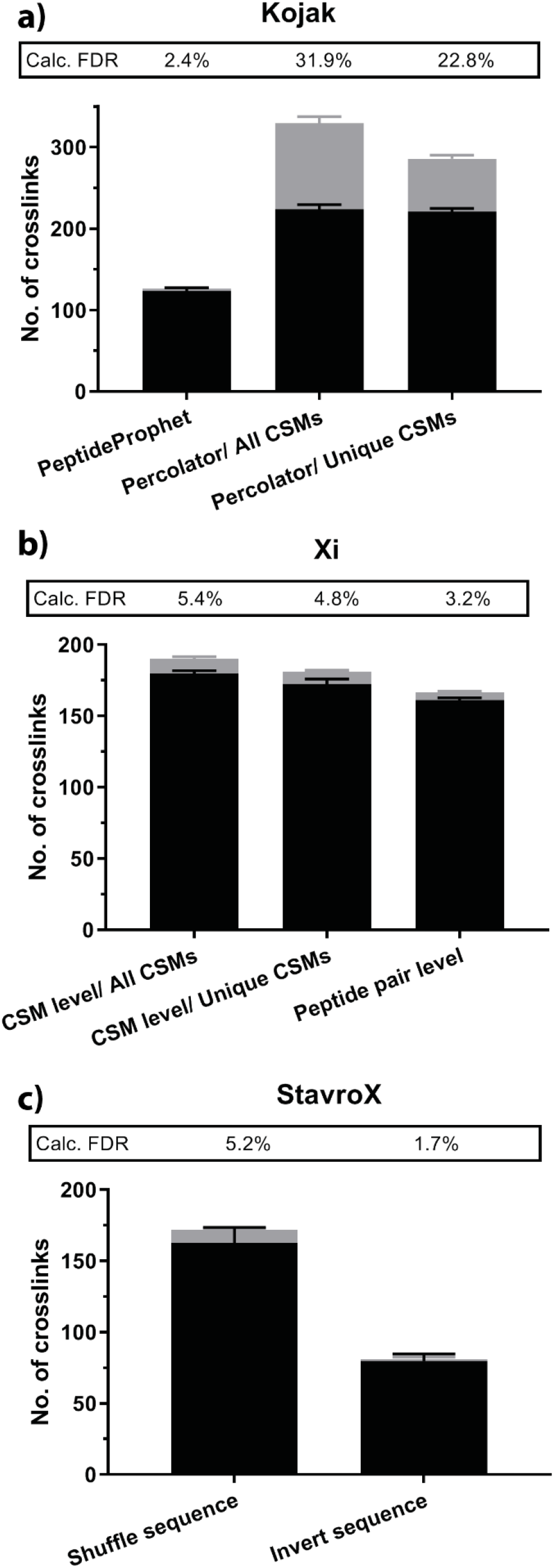
Effect of crosslink validation strategies on overall results. (a) Kojak results were validated by PeptideProphet, or Percolator with which all CSMs were used, or only the top-scoring CSM for each species. (b) Xi results were validated at the CSM level using all CSMs, the top-scoring CSM for each species or at the peptide pair level. (c) StavroX results were validated against a decoy database in which the sequences were shuffled with the protease sites remaining unchanged, or in which the sequences were inverted. All values are provided in Supplementary Table 6.

Fischer et al. [9] report that FDRs can be estimated at different points during data analysis, and it is implemented into Xi that the FDR can be calculated on different levels. We compared FDR estimations at the CSM level, the unique CSM level and the peptide pair level (Figure 3b). We note a slight decrease in the number of crosslinks identified when the FDR calculation is performed later in the analysis (180, 174 and 160), along with a decrease in calculated FDR (5.4%, 4.8% and 3.2%) for the CSM level, unique CSM level and peptide pair level, respectively.

StavroX offers two possibilities for generating the decoy database; to shuffle sequences and keep protease sites, or to invert the sequences. Crosslink validation with an inverted sequence is much more stringent than with the ‘shuffled sequence/ same protease sites’ option. While 163 crosslinks are found with the former (FDR 5.2%), just 80 crosslinks are identified with the latter, with an FDR of 1.7% which is much lower than the estimated cut-off of 5%.

We were also interested to see how the various search algorithms perform when the data is searched against a larger database (Supplementary Figure 2). For this we used a database containing the sequence of *S. pyogenes* Cas9 and 116 contaminant proteins (the crapome [27]). For pLink and the Kojak/Percolator combination, the larger database allowed a more accurate FDR estimation, along with a slight reduction in the number of correct crosslinks being identified. For pLink the FDR is reduced from 11.3% to 6.1% and the number of correct crosslinks is reduced from 217 to 204, while for Kojak/Percolator the FDR is reduced from 23% to 12% with a reduction in correct crosslinks from 221 to 202. Interestingly, the larger database had the opposite effect on the Xi results. With all FDR methods there is an increase in the number of correctly identified crosslinks (e.g. from 160 to 175 for the unique peptide level) along with an increase in calculated FDR (e.g. 3.2% to 6.7% for the unique peptide level). For the Kojak/PeptideProphet combination, the larger database increases the FDR (2.4% to 3.7%) while reducing the number of correctly identified crosslinks (123 to 121), both of which are undesirable effects in a crosslinking study.

### Comparison of scores attributed to CSMs

Another possible reason for the discrepancy between algorithms is the scoring functions. To investigate this further, scores attributed to CSMs by the various algorithms are compared (Figure 4). pLink scores given to each CSM are compared with those given by Xi (Figure 4a), StavroX (Figure 4b) and Kojak (Figure 4d), and scores given by Xi are compared to those given by StavroX (Figure 4c). CSMs correlating to correct crosslinks are shown in black, and CSMs correlating to false positives are shown in red. The scores attributed by pLink to false positive identifications are at the lower end of the scoring scale, suggesting that the pLink scoring system is robust. It is therefore expected that a slightly more stringent scoring approach would remove the false positives with little effect on the true crosslinks. There is positive correlation between the scores of pLink and Xi (R^2^=0.41) and the scores of StavroX and Xi (R^2^= 0.44), with lower correlation between pLink and StavroX (R^2^=0.15) and even lower between pLink and Kojak (R^2^= 0.06). This demonstrates that all algorithms have very different approaches for scoring crosslink assignments.

**Figure 4;.**
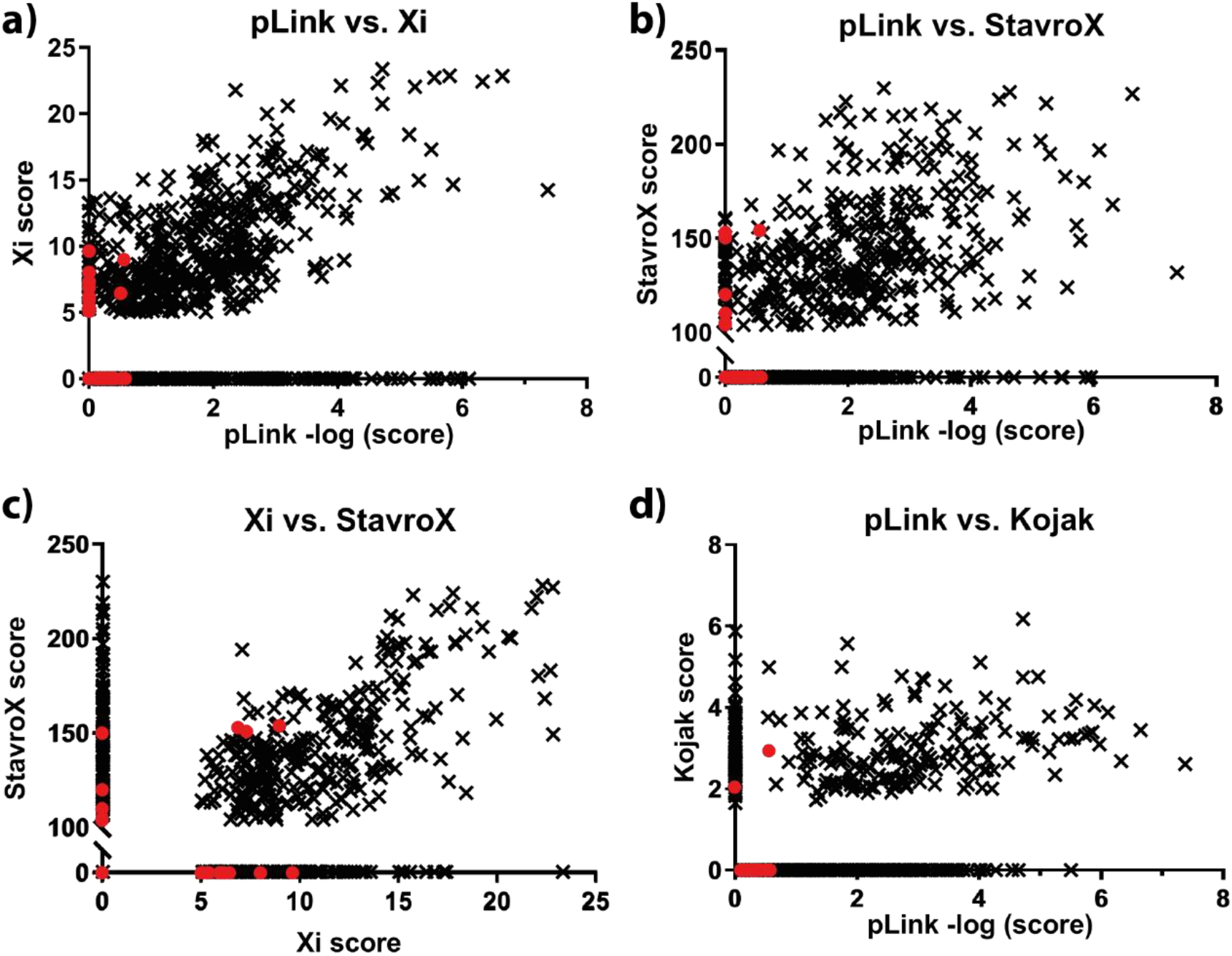
Comparison of scores assigned to each CSM by pLink, StavroX, Xi and Kojak. CSMs correlating to correct crosslinks are shown in black, and CSMs correlating to false positives are shown in red.

### Score distributions for candidate and decoy CSMs

The score distributions for the candidate and decoy peptides are shown for pLink (Figure 5a), Xi (Figure 5b) and StavroX (Figure 5c). The distribution of the scores given to decoy crosslinks correlates well with the stringency and accuracy of the algorithms. pLink attributes very low scores to the decoy crosslinks, and the score cut-off for the candidate crosslinks is therefore very low. StavroX on the other hand attributes a range of scores to decoy peptides, and the cut-off value for candidate crosslinks is very high, providing a stringent validation. For Xi, the situation lies between the two extremes; the number of correct and incorrect crosslinks lies between the number found for pLink and StavroX.

**Figure 5;.**
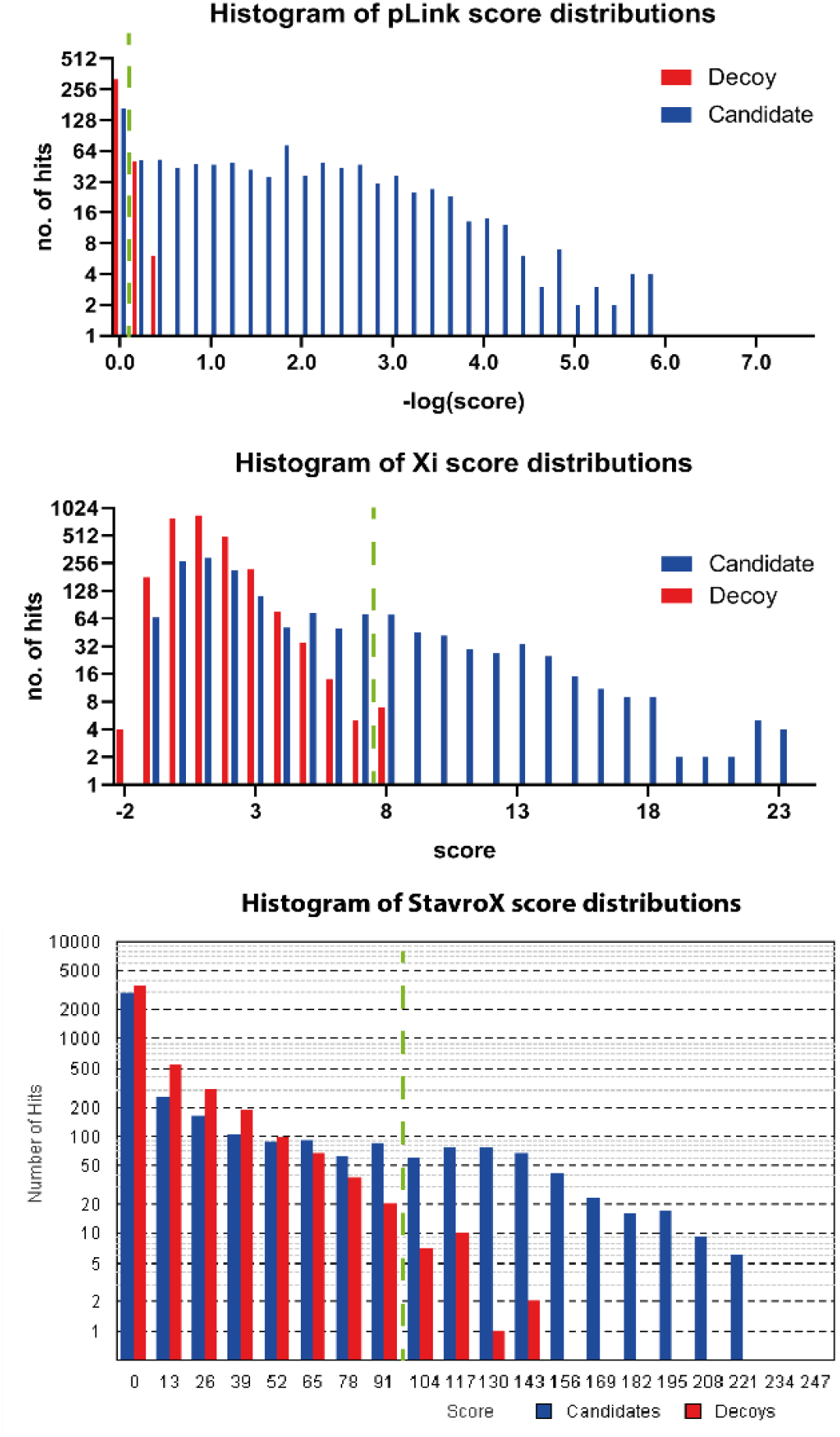
Score distributions for candidate and decoy crosslinks given by (a) pLink, (b) StavroX and (c) Xi. Here, the true positive and false positive crosslinks are grouped as candidates. Green dashed line refers to the score cut-off used to filter for an estimated 5% FDR.

### Number of MS2 spectra utilised by the search engines

During measurement of the first technical replicate of the DSS-crosslinked peptides, 5022 MS2 spectra were triggered during the MS acquisition. In the case of pLink, 666 of these were assigned to crosslinks, 172 were assigned to monolinks, looplinks and unmodified peptides, 168 were assigned to candidate crosslinks that had a score below the cut-off for the estimated 5% FDR, 383 were assigned to decoy PSMs (corresponding to all species) and 3633 triggered spectra were not utilised in the search (Figure 6). From the 168 spectra that had a score below the cut-off value for 5% FDR, just 4 additional correct crosslinks were identified that failed to pass the validation. An additional 105 incorrect crosslinks were found in this ‘unvalidated’ portion of the CSMs, demonstrating efficient separation of crosslinks that are correctly and incorrectly assigned.

**Figure 6;.**
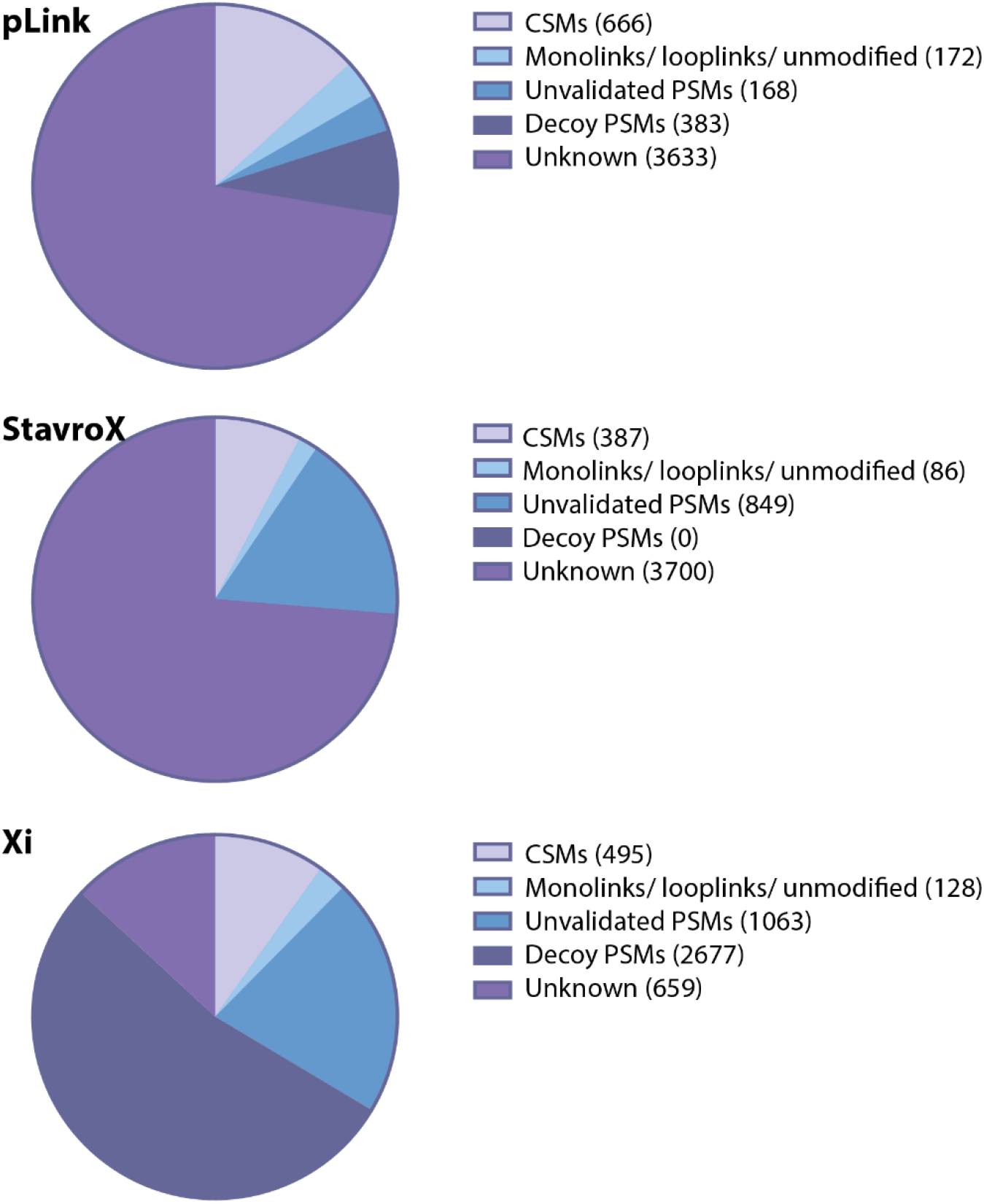
number of MS2 spectra that are attributed to (i) CSMs, (ii) monolinks/ looplinks or unmodified peptides, (iii) PSMs that have a score below the cut-off value for 5% FDR, (iv) decoy PSMs and (v) those that are unused by the algorithm.

StavroX doesn’t provide information on unmodified peptides, or the spectra that were assigned to decoy crosslinks. Nevertheless, a much higher number of spectra were assigned to unvalidated species that had a score below the cut-off for an estimated 5% FDR than pLink. From these, an additional 58 correct crosslinks were identified, and 389 incorrect. This suggests that StavroX doesn’t have a problem with identifying crosslinks, but rather has difficulty distinguishing those that are correct from those that are incorrect.

Xi ascribes 2677 spectra to decoy species, much higher than pLink (383), which is a potential reason for it being a more stringent search algorithm. 11 additional correct crosslinks can be identified in the ‘non-validated’ CSMs, which is a much lower number than for StavroX, while 757 additional incorrect crosslinks are identified, which is more than both StavroX and pLink. Xi therefore appears to have a robust method of separating correct crosslinks from incorrect.

### Assessment of workflows developed for cleavable crosslinkers

Recently, MS-cleavable crosslinkers have been developed to increase confidence of crosslink assignments and to allow identification of crosslinks in more complex samples. Here, the peptides were crosslinked with DSBU or DSSO and measured using stepped HCD on an Orbitrap Q-Exactive HF-X (Figure 7). Data are searched against a database containing the sequence of *S. pyogenes* Cas9 and the crapome [27] with MeroX 2.0 (in Rise and RiseUP mode) [28] and XlinkX in Proteome Discoverer 2.3 [29]. This larger database was used for analysis of peptides crosslinked with the cleavable reagents compared to DSS, because a key benefit of such crosslinkers is the ability to search data against larger databases, up to the scale of the human proteome. Also, it has been recommended by the developers to use a database with at least 100 other protein sequences that are used as support for FDR control [29].

**Figure 7;.**
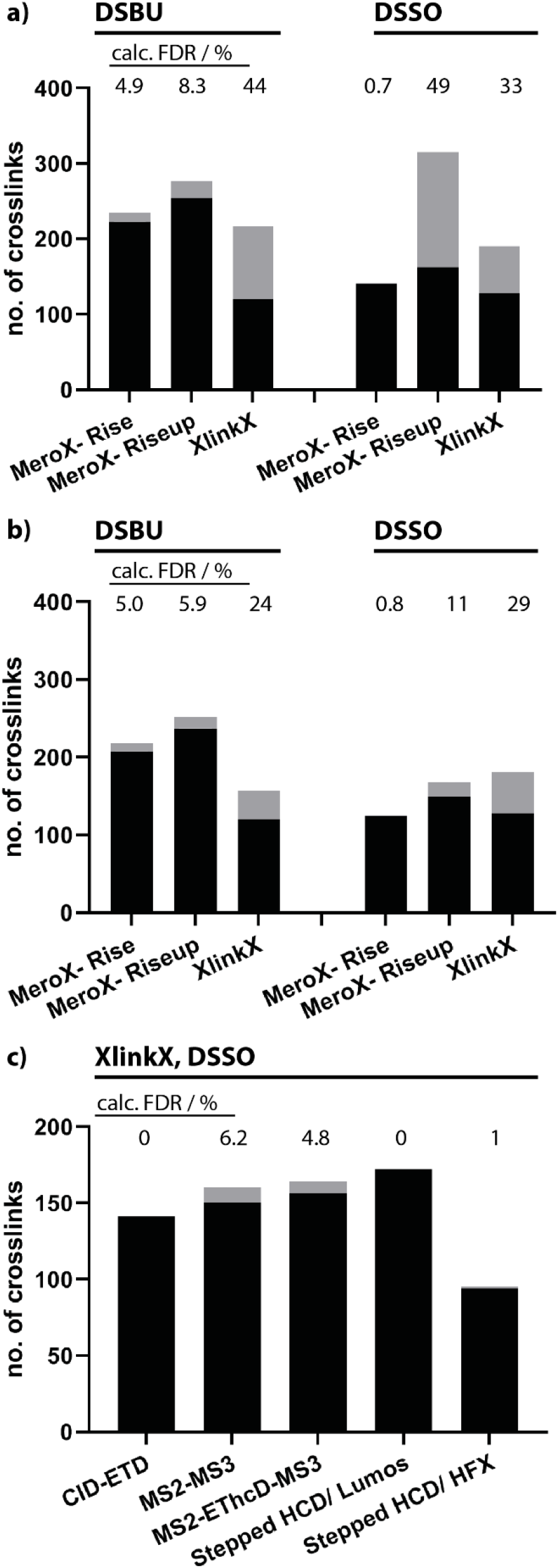
Assessment of algorithms developed for the identification of peptides crosslinked with cleavable reagents DSBU and DSSO. (a, b) Data were collected using MS2 method with stepped collision energy, analysed using MeroX (in Rise and RiseUP mode) and XlinkX, with data filtered to an estimated 5% FDR (a) and 1% FDR (b), with no additional score cut-off values employed. (c) Data for DSSO-crosslinked peptides were collected using HCD- MS3- and ETD-based methods, and analysed using XlinkX with the recommended score cut-off values of 45 and 4 for crosslink score and Δcrosslink score, respectively. Values are given in Supplementary Tables 8-10.

We first analysed data recorded on a Q-Exactive HFX with a stepped HCD fragmentation method as recommended by Iacobucci et al [28]. Data were filtered to an estimated FDR of 5% with no additional score cut-off values. When data from the measurement of DSBU peptides are analysed by MeroX in Rise mode (Figure 7a), in which 3 out of 4 reporter peaks must be present in the MS2 spectrum, 223 correct crosslinks and 12 incorrect crosslinks are identified, giving rise to a calculated FDR of 4.9%. When used in RiseUP mode, which appears to be a less stringent analysis method, the number of correct crosslinks increases to 254 but the number of false identifications also rises to 23 resulting in an increased FDR of 8.3%. XlinkX finds 120 correct crosslinks and 96 false positives, resulting in an FDR of 44%. One reason that fewer crosslinks are detected with XlinkX is that, unlike MeroX, it doesn’t report crosslinks between two peptides of the same sequence. When such crosslinks are removed from the MeroX search results, the number of correctly identified crosslinks are 169 and 199 for Rise and RiseUP mode, respectively. MeroX identifies a lower number of DSSO crosslinked peptides than DSBU, likely because MeroX was developed for the analysis of DSBU crosslinked samples. In Rise mode 140 true crosslinks are identified (with 1 false positive), and in RiseUP mode 162 true crosslinks are identified (153 false positives). XlinkX identifies 128 true DSSO crosslinks (with 62 false positives), which is 8 more than were identified with DSBU.

When the estimated FDR cut-off was reduced to 1%, the FDR for the DSBU/Rise combination actually increased to 5%, the DSBU/Riseup combination reduced to 5.9% and the DSBU/XlinkX combination reduced to 24%. For the DSSO crosslinked peptides the FDR is reduced to 0.8% for Rise, 11% for Riseup and 29% for XlinkX. We were surprised to note from this study that the estimated FDR is often less reliable for cleavable crosslinkers than can be achieved with DSS (Figure 2).

Methods involving MS3 and ETD that were developed for the measurement of DSSO-crosslinked peptides on an Orbitrap Fusion Lumos were also investigated, and the data were analysed using XlinkX (Figure 7c). Here, we also employed the recommended score cut-off of 45 and minimum score difference of 4 [29]. Upon measurement with CID-ETD, 141 crosslinks were identified, all of which were correct. MS2-MS3 measurements allowed identification of 150 correct and 10 incorrect crosslinks, resulting in an FDR of 6.2%. The hybrid method MS2-EThcD-MS3 gives rise to 156 correct and 8 incorrect crosslinks (calculated FDR 4.9%). Measurement with stepped HCD gives rise to the highest number of true crosslinks (172) with just one false positive (0.5% FDR). When the data from the Q-exactive HFX were analysed with these score cut-offs 94 correct and 1 incorrect crosslinks were identified (1.1% FDR).

## Discussion

This unique synthetic crosslinked peptide library is an excellent resource for the crosslinking community. The data arising from analysis of the crosslinked peptides can be used for in-depth assessment of the many algorithms available for XL identification. This provides valuable information for users of XL-MS regarding the performance of crosslink search algorithms, allowing them to select data analysis strategies that meets their needs in terms of confidence and sensitivity of crosslink assignments. The data will also be invaluable in the development of crosslink search engines, since, for the first time, it is known which crosslinks are true, which are false, and which remain to be identified. This allows better assessment of different FDR calculations that are currently available.

This study also allows assertions to be made about the analysis of crosslinking data. Eminently, when using pLink, or the Kojak/Percolator combination, a large database is required for better FDR estimation. A reason for better performance with the larger database could be the higher number of CSMs in the resulting set that increase the training size for the algorithm, or it could be that false CSMs are less likely to be randomly assigned to Cas9, thereby producing a more defined decoy population exploited by the algorithms during the training regimen. This is particularly likely to be true for Percolator, that performs best when trained on 100,000 or more spectra [18], which is far more than the 5022 contained in this dataset. Alternatively, Xi and Kojak/PeptideProphet perform better with a smaller dataset, for example with 10 proteins. The data included in this publication can be utilised to further optimise such parameters.

Additionally, when using XlinkX for the analysis of data in which cleavable crosslinkers were used, the score cut-offs are important in filtering for high-quality data that more likely allow correct crosslink identification. One should be aware, that while such cut-offs work well in this study and others [30, 31], they give no indication of the confidence in crosslink assignment. Judging by the data, different cut-off values could be optimised for different fragmentation techniques. For example, a lower value of 20 could be used for the stHCD data that gives rise to 183 correct crosslinks (11 more than with a score cut-off of 40) and 8 incorrect, resulting in a FDR of 4.1%. A similar approach was taken by Ser et al. [7] by spiking crosslinked BSA peptides into non-crosslinked proteome background peptides to calculated score filters that removed 99% and 90% of non-BSA crosslinks to determine FDR cut-offs for 1% and 10% FDR, respectively. The data included in the present publication could be even more useful for a systematic optimisation of search and validation strategies.

Results from XL-MS studies have far-reaching implications, often leading to new avenues of research regarding novel protein-protein interactions. It is therefore of paramount importance that the confidence of identified crosslinks is correctly estimated. This peptide library is the first true assessment of crosslink confidence. As well as providing a useful bioinformatics resource, the physical library can be used to optimise many stages of the crosslinking workflow including crosslink enrichment strategies, chromatography methods and MS data acquisition parameters.

## Methods

### Peptide design and synthesis

Peptides were synthesised on a SYRO with Tip Synthesis Module (MultiSynTech GmbH) using standard Fmoc chemistry. For each amino acid cycle, double coupling with DIC/K-Oxyma and HATU/DIEA was performed. C-terminal lysine residues were initially incorporated with an azide group. Peptides were purified on a C18 kinetex column (2.6 μm) using a 30 minute gradient. Peptide concentration was measured using a nanodrop, the solution was evaporated to almost dryness, and the peptides were resuspended at a concentration of 5mM in HEPES buffer (100mM pH 8) and the appropriate peptides were combined into groups (Supplementary Table 1).

### Crosslinking

Crosslinker was dissolved at 20mM in DMSO, and 0.5 μl were added to 5 μl to each peptide group 5x over the course of 2.5 hours. 1 μl of each crosslinked peptide group was added to 9 μl ammonium bicarbonate (ABC, 100 mM) for overnight digestion with 10ng trypsin at 37 °C and subsequent reduction with tris(2-carboxyethyl)phosphine for 30 minutes (final concentration 50 mM). All 12 groups were combined into one sample which was stored at −80° C in 5 μl aliquots for future use.

### Biotin removal

After digestion, the biotin-linker region was removed with streptavidin beads. 25ul of slurry was washed 3 times with 25ul ABC buffer. 5ul peptides + 20ul ABC buffer was loaded onto the beads and incubated for 30 minutes. The eluate was removed and the beads were washed with 25ul 0.1% FA and subsequently 25ul 30% MeOH. The eluates were combined and reduced to 5ul in a centrifugal concentrator to remove the MeOH. The sample was then diluted 5x in 0.1% TFA and 1ul of the final solution was injected.

### Reversed-phase high-performance liquid chromatography

crosslinked peptides were separated using a Dionex UltiMate 3000 high-performance liquid chromatography (HPLC) RSLCnano System prior to MS analysis. The HPLC was interfaced with the mass spectrometer via a Nanospray Flex ion source. For sample concentrating, washing, and desalting, the peptides were trapped on an Acclaim PepMap C-18 precolumn (0.3 × 5 mm, Thermo Fisher Scientific) using a flow rate of 25 μL/min and 100% buffer A (99.9% H2O, 0.1% TFA). The separation was performed on an Acclaim PepMap C-18 column (50 cm × 75 μm, 2 μm particles, 100 Å pore size, Thermo Fisher Scientific) applying a flow rate of 230 nL/min. For separation, a solvent gradient ranging from 2 to 40% buffer B (80% ACN, 19.92% H2O, 0.08% TFA) was applied. The gradient length was 2 hours for the analysis of the combined groups.

### Mass spectrometry

MS settings were used according to previously published protocols. DSS-, DSSO- and DSBU-crosslinked peptides were measured on both an Orbitrap Q-Exactive HF-X instrument and an Orbitrap Fusion Lumos mass spectrometer. DSS samples were measured with the settings recommended by Chen and Rappsilber [16]. DSBU measurements on the HFX were measured with the settings recommended by Iacobucci et al. [28]. DSSO samples were measured on an Orbitrap Fusion Lumos mass spectrometer with the settings recommended by Klykov et al. [29] for MS3 and ETD-based acquisitions, and those recommended by Stieger et al. [32] for stepped HCD methods. DSSO samples were measured on the HFX in data-dependent MS/MS mode with stepped HCD and normalised collision energies of 21, 27 and 33 %. Each high-resolution full scan (m/z 375-1500, R = 60 000) in the orbitrap is followed by ten high-resolution production scans (R = 30 000), starting with the most intense signal in the full scan mass spectrum (isolation window 1.6 m/z). The target value of the automated gain control is set to 5e5 (MS) and 5e4 (MS/MS) and maximum accumulation times are set to 50 ms (MS) and 100 ms (MS/MS). Precursor ions with charge states < 3+ or >8+ are excluded from fragmentation. Dynamic exclusion is enabled (duration 30 s; window 10 ppm).

### Data analysis

Raw data collected from the measurement of DSS-crosslinked peptides were converted to .mgf format using MSconvert with the Peak Picking function and searched with the relevant programmes against a database containing *S. pyogenes* Cas9 and 10 additional proteins. Settings for each programme are given in Supplementary Information Table 11. Data collected from the measurement of cleavable crosslinkers were searched with MeroX and XlinkX against a database containing *S. pyogenes* Cas9 and the Crapome [27]. Settings are given in Supplementary Information Table 12.

### Data Availability

The mass spectrometry proteomics data have been deposited to the ProteomeXchange Consortium via the PRIDE [33] partner repository with the dataset identifier PXD014337. Username: Password: DDNDTsZK

Results from each search engine are available as .CSV files upon request.

## Acknowledgements

The authors acknowledge Andre Luttig and Gerhard Dürnberger for help with data analysis, and all members of the protein chemistry facility for fruitful discussions. The authors acknowledge Mathias Madalinski for synthesising the peptides. RB acknowledges the Austrian Science Fund for the receipt of a Lise Meitner Postdoctoral Fellowship (project number M2334). JS is a Wittgenstein Prize Fellow and funded by the T. von Zastrow Foundation. J.M.P. is supported by grants from IMBA, the Austrian Academy of Sciences, the T. von Zastrow Foundation, an ERC Advanced Grant, and an Era of Hope Innovator award. KM has been supported by EPIC-XS, project number 823839, funded by the Horizon 2020 program of the European Union, and the Austrian Science Fund by ERA-CAPS I 3686 International Project.

## Author Contributions

R.B and J.S. planned the experiments, and R.B. carried them out. R.B. analysed the data and wrote the manuscript, which was edited by J.S. K.M. and J.M.P supervised the experiments.

## Competing interests

The authors declare no competing interests

## Supplementary information

Further details regarding crosslink search settings and results can be found in supplementary information.

## Notes

https://www.ebi.ac.uk/pride/archive/

