## supplementary information for "A synthetic peptide library for benchmarking crosslinking mass spectrometry search engines"

**Supplementary Table 1 - peptide sequences in each group.**

**Supplementary Table 2 - Number of CSMs attributed to DSS crosslinks by pLink, StavroX and Xi. Results were filtered to an estimated 5% FDR.**

**Supplementary Table 3 - Number of DSS crosslinks identified with pLink, StavroX and Xi. Results were filtered to an estimated 5% FDR.**

**Supplementary Figure 1 - Number of CSMs/ crosslinks identified by pLink, StavroX and Xi. Results were filtered to an estimated 1% FDR**

**Supplementary Table 4 - Number of CSMs attributed to DSS crosslinks by pLink, StavroX and Xi. Results were filtered to an estimated 1% FDR.**

**Supplementary Table 5 - Number of DSS crosslinks identified with pLink, StavroX and Xi. Results were filtered to an estimated 1% FDR.**

**Supplementary Table 6 - Number of DSS crosslinks identified with Kojak, Xi and StavroX employing different validation strategies, as shown in Figure 3.**

**Supplementary Figure 2 - Number of DSS crosslinks identified when the data were searched against the Cas9 sequence and the CrapDB**

**Supplementary Table 7 - Number of DSS crosslinks identified when the data were searched against the Cas9 sequence and the CrapDB**

**Supplementary Table 8 - Number of DSBU or DSSO crosslinks identified with MeroX or XlinkX upon analysis of data generated with stepped HCD on a Q-exactive HFX instrument. Results were filtered to an estimated 5% FDR**

**Supplementary Table 9 - Number of DSBU or DSSO crosslinks identified with MeroX or XlinkX upon analysis of data generated with stepped HCD on a Q-exactive HFX instrument. Results were filtered to an estimated 1% FDR**

**Supplementary Table 10; Number of crosslinks identified with XlinkX upon analysis of the data generated with different fragmentation strategies.**

**Supplementary Table 11 - Search settings used for the identification of DSS- crosslinked peptides.**

**Supplementary Table 12; Search settings used for the identification of DSBU- and DSSO- crosslinked peptides.**

**Sequences used during the database search of DSS-crosslinked peptides.**

| Group | Peptide sequences |
|---|---|
| **Group 1** | SDKNR |
| | KLINGIR |
| | KFDNLTK |
| | FIKPILEK |
| | APLSASMIKR |
| | NPIDFLEAKGYK |
| | LPKYSLFELENGR |
| | TEVQTGGFSKESILPK |
| **Group 2** | VKYVTEGMR |
| | FDNLTKAER |
| | DFQFYKVR |
| | YDENDKLIR |
| | MIAKSEQEIGK |
| | HKPENIVIEMAR |
| | TILDFLKSDGFANR |
| | KIECFDSVEISGVEDR |
| | YVNFLYLASHYEKLK |
| **Group 3** | LSKSR |
| | DKPIR |
| | KDLIIK |
| | MKNYWR |
| | KGILQTVK |
| | NSDKLIAR |
| | DDSIDNKVLTR |
| **Group 4** | KLVDSTDK |
| | IEKILTFR |
| | KAIVDLLFK |
| | VLSAYNKHR |
| | IEEGIKELGSQILK |
| | SSFEKNPIDFLEAK |
| | SNFDLAEDAKLQLSK |
| | HSLLYEYFTVYNELTKVK |
| **Group 5** | KVTVK |
| | EKIEK |
| | VITLKSK |
| | QLKEDYFK |
| | QLLNAKLITQR |
| | GGLSELDKAGFIK |
| | MDGTEELLVKLNR |
| **Group 6** | EVKVITLK |
| | KPAFLSGEQK |
| | ENQTTQKGQK |
| | KTEVQTGGFSK |
| | VVDELVKVMGR |
| | LESEFVYGDYKVYDVR |
| | MLASAGELQKGNELALPSK |
| | NFMQLIHDDSLTFKEDIQK |
| | VLPKHSLLYEYFTVYNELTK |
| **Group 7** | KMIAK |
| | ESILPKR |
| | DLIIKLPK |

| | |
|---|---|
| | FKVLGNTDR<br>SEQEIGKATAK<br>AIVDLLFKTNR<br>LKTYAHLFDDK<br>VNTEITKAPLSASMIK<br>YDEHHQDLTLLKALVR |
| **Group 8** | KDWDPK<br>QQLPEKYK<br>KVLSMPQVNIVK<br>MTNFDKNLPNEK<br>QITKHVAQILDSR<br>KSEETITPWNFEEVVDK<br>KNGLFGNLIALSLGLTPNFK<br>SKLVSDFR |
| **Group 9** | LKSVK<br>IIKDK<br>DWDPKK<br>LKGSPEDNEQK<br>VLSMPQVNIVKK<br>LENLIAQLPGEKK<br>LIYLALAHMIKFR<br>YPKLESEFVYGDYK |
| **Group 10** | VPSKK<br>VTVKQLK<br>EDYFKK<br>VKYVTEGMR<br>GKSDNVPSEEVVK<br>LEESFLVEEDKK<br>QEDFYPFLKDNR |
| **Group 11** | GQKNSR<br>AGFIKR<br>GYKEVK<br>VMKQLK<br>KDFQFYK<br>LVDSTDKADLR<br>SDNVPSEEVVKK<br>KNLIGALLFDSGETAEATR |
| **Group 12** | HSIKK<br>DKQSGK<br>NLPNEKVLPK<br>QSGKTILDFLK<br>MNTKYDENDK<br>SVKELLGITIMER<br>TYAHLFDDKVMK<br>FNASLGTYHDLLKIIK |

**Supplementary Table 1; peptide sequences in each group.**

| Search engine | Number of crosslink- spectrum matches | | | | | | | | |
|---|---|---|---|---|---|---|---|---|---|
| | Correct | | | Incorrect | | | Calculated FDR (%) | | |
| | R1 | R2 | R3 | R1 | R2 | R3 | R1 | R2 | R3 |
| **pLink** | 639 | 712 | 683 | 27 | 27 | 39 | 4.1 | 3.7 | 5.4 |
| **StavroX** | 378 | 434 | 419 | 9 | 12 | 10 | 2.3 | 2.7 | 2.3 |
| **Xi** | 491 | 498 | 547 | 20 | 13 | 10 | 3.9 | 2.6 | 1.8 |

**Supplementary Table 2;** Number of CSMs attributed to DSS crosslinks by pLink, StavroX and Xi. Results were filtered to an estimated 5% FDR. Measurements were performed in technical triplicate (R1, R2, R3).

| Search engine | Number of crosslinks | | | | | | | | |
|---|---|---|---|---|---|---|---|---|---|
| | Correct | | | Incorrect | | | Calculated FDR (%) | | |
| | R1 | R2 | R3 | R1 | R2 | R3 | R1 | R2 | R3 |
| **pLink** | 217 | 230 | 203 | 26 | 24 | 33 | 10.7 | 9.4 | 14.0 |
| **StavroX** | 159 | 175 | 154 | 8 | 10 | 9 | 4.8 | 5.4 | 5.5 |
| **Xi** | 179 | 183 | 179 | 18 | 11 | 7 | 9.1 | 5.7 | 3.8 |

**Supplementary Table 3;** Number of DSS crosslinks identified with pLink, StavroX and Xi. Results were filtered to an estimated 5% FDR. Measurements were performed in technical triplicate (R1, R2, R3).

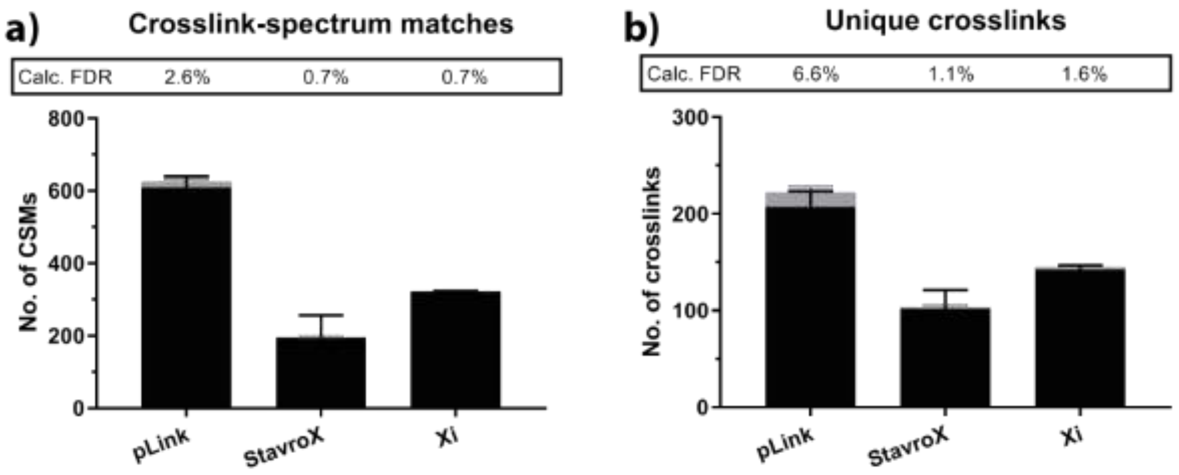

**Supplementary Figure 1.** (a) number of CSMs that correspond to correct (black) and incorrect (grey) crosslinks identified by pLink, StavroX and Xi. Results were filtered to an estimated 1% FDR, and the calculated FDR is given for each algorithm. Error bars correspond to the standard deviation between three technical replicates. (b) number of correct unique crosslinks (black) and incorrect crosslinks (grey). Values are given in Supplementary tables 4 and 5.

| Search engine | Number of crosslink- spectrum matches | | | | | | | | |
|---|---|---|---|---|---|---|---|---|---|
| | Correct | | | Incorrect | | | Calculated FDR (%) | | |
| | R1 | R2 | R3 | R1 | R2 | R3 | R1 | R2 | R3 |
| pLink | 594 | 644 | 585 | 10 | 13 | 25 | 1.7 | 2.0 | 4.1 |
| StavroX | 265 | 157 | 160 | 4 | 0 | 1 | 1.5 | 0 | 0.6 |
| Xi | 312 | 352 | 438 | 2 | 4 | 5 | 0.6 | 1.1 | 1.1 |

**Supplementary Table 4;** Number of CSMs attributed to DSS crosslinks by pLink, StavroX and Xi. Results were filtered to an estimated 1% FDR. Measurements were performed in technical triplicate (R1, R2, R3).

| Search engine | Number of crosslinks | | | | | | | | |
|---|---|---|---|---|---|---|---|---|---|
| | Correct | | | Incorrect | | | Calculated FDR (%) | | |
| | R1 | R2 | R3 | R1 | R2 | R3 | R1 | R2 | R3 |
| **pLink** | 215 | 218 | 189 | 9 | 12 | 22 | 4.0 | 5.2 | 11.6 |
| **StavroX** | 124 | 91 | 90 | 4 | 0 | 1 | 3.1 | 0 | 1.1 |
| **Xi** | 141 | 152 | 163 | 2 | 3 | 5 | 1.4 | 1.9 | 3.0 |

**Supplementary Table 5;** Number of DSS crosslinks identified with pLink, StavroX and Xi. Results were filtered to an estimated 1% FDR. Measurements were performed in technical triplicate (R1, R2, R3).

| Search engine | Number of crosslinks | | | | | | | | |
|---|---|---|---|---|---|---|---|---|---|
| | Correct | | | Incorrect | | | Calculated FDR (%) | | |
| | R1 | R2 | R3 | R1 | R2 | R3 | R1 | R2 | R3 |
| Kjk, PepProphet | 128 | 121 | 120 | 2 | 3 | 4 | 1.5 | 2.4 | 3.2 |
| Kjk, Perc, all | 222 | 230 | 219 | 96 | 108 | 112 | 30.1 | 32.0 | 33.8 |
| Kjk, Perc, unique | 220 | 225 | 217 | 68 | 60 | 67 | 23.6 | 21.0 | 23.6 |
| Xi, all CSMs | 179 | 183 | 179 | 18 | 11 | 7 | 9.1 | 5.7 | 3.8 |
| Xi, unique CSMs | 176 | 175 | 170 | 6 | 10 | 6 | 3.3 | 5.4 | 3.4 |
| Xi, peptide pair | 162 | 161 | 158 | 6 | 4 | 5 | 3.6 | 2.4 | 3.1 |
| StavroX, Shuffle | 159 | 175 | 154 | 8 | 10 | 9 | 4.8 | 5.4 | 5.5 |
| StavroX, invert | 74 | 81 | 84 | 3 | 0 | 1 | 3.9 | 0 | 1.2 |

**Supplementary Table 6;** Number of DSS crosslinks identified with Kojak, Xi and StavroX employing different validation strategies, as shown in Figure 3. Results are filtered to an estimated 5% FDR.

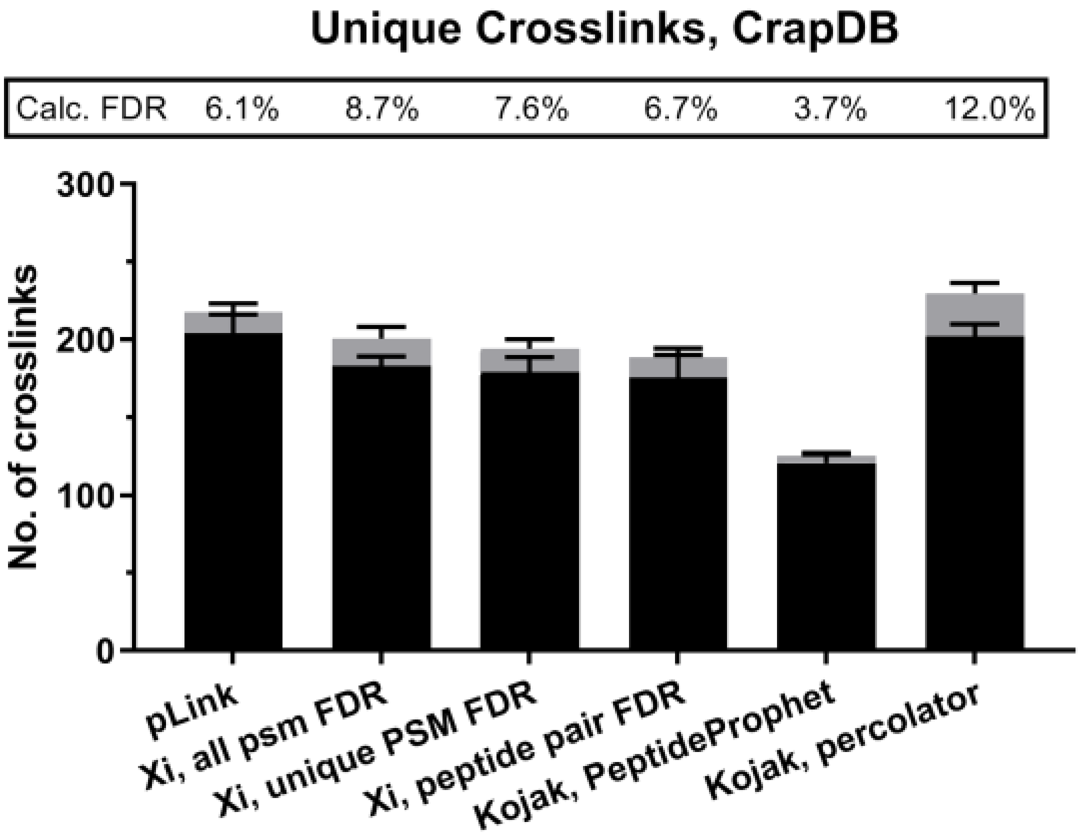

**Supplementary Figure 2** Number of correct (black) and incorrect (grey) unique crosslinks identified when the data were searched against the Cas9 sequence and the CrapDB, which contains 116 proteins.

| Search engine | Number of crosslinks | | | | | | | | |
|---|---|---|---|---|---|---|---|---|---|
| | Correct | | | Incorrect | | | Calculated FDR (%) | | |
| | **R1** | **R2** | **R3** | **R1** | **R2** | **R3** | **R1** | **R2** | **R3** |
| **pLink** | 215 | 206 | 191 | 18 | 7 | 15 | 7.7 | 3.3 | 7.3 |
| **Xi, all CSM** | 181 | 190 | 178 | 22 | 22 | 9 | 10.8 | 10.4 | 4.8 |
| **Xi, unique CSM** | 180 | 188 | 169 | 19 | 18 | 8 | 9.5 | 8.7 | 4.5 |
| **Xi, peptide pair** | 179 | 188 | 159 | 17 | 16 | 6 | 8.7 | 7.8 | 3.6 |
| **Kojak, PepProph** | 126 | 113 | 123 | 4 | 4 | 6 | 3.1 | 3.4 | 4.7 |
| **Kojak, Perc (unique)** | 198 | 211 | 197 | 20 | 31 | 32 | 9.1 | 12.8 | 14.0 |

**Supplementary Table 7;** number of DSS crosslinks identified with pLink, Xi and Kojak when data were searched against the CrapDB, containing 116 proteins.

| MS method | Number of crosslinks | | |
|---|---|---|---|
| | Correct | Incorrect | Calculated FDR (%) |
| **DSBU, MeroX, Rise** | 223 | 12 | 4.9 |
| **DSBU, MeroX, Riseup** | 254 | 23 | 8.3 |
| **DSBU, XlinkX** | 120 | 96 | 44 |
| **DSSO, MeroX, Rise** | 140 | 1 | 0.7 |
| **DSSO, MeroX, Riseup** | 162 | 153 | 49 |
| **DSSO, XlinkX** | 128 | 62 | 33 |

**Supplementary Table 8;** Number of DSBU or DSSO crosslinks identified with MeroX or XlinkX upon analysis of data generated with stepped HCD on a Q-exactive HFX instrument. Results were filtered to an estimated **5% FDR** with no extra score cut-offs implemented.

| MS method | Number of crosslinks | | |
|---|---|---|---|
| | Correct | Incorrect | Calculated FDR (%) |
| **DSBU, MeroX, Rise** | 207 | 11 | 5.0 |
| **DSBU, MeroX, Riseup** | 237 | 15 | 5.9 |
| **DSBU, XlinkX** | 120 | 37 | 24 |
| **DSSO, MeroX, Rise** | 124 | 1 | 0.8 |
| **DSSO, MeroX, Riseup** | 149 | 19 | 11 |
| **DSSO, XlinkX** | 128 | 53 | 29 |

**Supplementary Table 9;** Number of DSBU or DSSO crosslinks identified with MeroX or XlinkX upon analysis of data generated with stepped HCD on a Q-exactive HFX instrument. Results were filtered to an estimated **1% FDR** with no extra score cut-offs implemented.

| MS method | Number of crosslinks | | |
|---|---|---|---|
| | Correct | Incorrect | Calculated FDR (%) |
| CID-ETD | 141 | 0 | 0 |
| MS2-MS3 | 150 | 10 | 6.2 |
| MS2-EThcD-MS3 | 156 | 8 | 4.8 |
| Stepped HCD, Lumos | 172 | 0 | 0 |
| Stepped HCD, HFX | 94 | 1 | 1 |

**Supplementary Table 10;** Number of crosslinks identified with XlinkX upon analysis of the data generated with different fragmentation strategies. Score cut-off values were implemented of 45 and 4 for crosslink score and Δcrosslink score, respectively.

| | pLink (2.3.5) | StavroX (3.6.0) | Xi (1.6.751) | Kojak (1.6.1) |
|---|---|---|---|---|
| **Crosslink mass/ Da** | 138.068 | 138.068 | 138.068 | 138.068 |
| **Monolink mass/ Da** | 156.079 | 156.079 | 156.079 | 156.079 |
| **Crosslinker reactivity** | K-K | K-K | K-K | K-K |
| **Fixed modification** | Carbamido-methyl [C] | Carbamido-methyl [C] | Carbamido-methyl [C] | Carbamido-methyl [C] |
| **Variable modification** | Oxidation [M] | Oxidation [M] | Oxidation [M] | Oxidation [M] |
| **Enzyme** | Trypsin | Trypsin | Trypsin | Trypsin |
| **Max. missed cleavages** | 3 | R:3 K:3 | 3 | 3 |
| **Min peptide mass** | 500 | 500 | - | 500 |
| **Max peptide mass** | 6000 | 6000 | - | 6000 |
| **Min peptide length** | 5 | 5 | 5 | - |
| **Max peptide length** | 60 | - | - | - |
| **MS1 tolerance (ppm)** | 5 | 5 | 5 | 5 |
| **MS2 tolerance (ppm)** | 20 | 20 | 20 | Bin size 0.03 Thomson |
| **FDR calculation** | inbuilt | inbuilt | Xi FDR (1.1.27) | Percolator (3.02) |
| **FDR level** | PSM | PSM | PSM | PSM |

**Supplementary Table 11;** Search settings used for the identification of DSS- crosslinked peptides.

| | XlinkX in proteome discoverer 2.3 | MeroX 2.0 beta 5 |
|---|---|---|
| **DSBU Crosslink mass/ Da** | 196.085 | 196.085 |
| **Bu-fragment/ Da** | - | 85.053 |
| **BuUr-fragment/ Da** | - | 111.032 |
| **DSSO crosslink mass/ Da** | 158.004 | 158.004 |
| **Alkene/ Da** | - | 54.011 (essential) |
| **Thiol/ Da** | - | 85.983 (essential) |
| **Sulfenic acid/ Da** | - | 103.993 |
| **Crosslinker reactivity** | K-K | K-K |
| **Fixed modification** | Carbamidomethyl [C] | Carbamidomethyl [C] |
| **Variable modification** | Oxidation [M] | Oxidation [M] |
| **Enzyme** | Trypsin | Trypsin |
| **Max. missed cleavages** | 3 | R:3 K:3 |
| **Min peptide mass** | 500 | 500 |
| **Max peptide mass** | 6000 | 6000 |
| **Min peptide length** | 5 | 5 |
| **MS1 tolerance (ppm)** | 5 | 5 |
| **MS2 tolerance (ppm)** | 20 | 20 |
| **S/N ratio** | 1.5 | 1.5 |
| **FDR calculation** | inbuilt | inbuilt |
| **FDR level** | PSM | PSM |

**Supplementary Table 12**; Search settings used for the identification of DSBU- and DSSO- crosslinked peptides.

**Sequences used during the database search of DSS-crosslinked peptides. 10 proteins were taken at random from the CRAPome.**

>Cas9
GAASMDKKYSIGLAIGTNSVGWAVITDEYKVPSKKFKVLGNTDRHSIKKNLIG
ALLFDSGETAEATRLKRTARRRYTRRKNRICYLQEIFSNEMAKVDDSFFHRLE
ESFLVEEDKKHERHPIFGNIVDEVAYHEKYPTIYHLRKKLVDSTDKADLRLIYL
ALAHMIKFRGHFLIEGDLNPDNSDVDKLFIQLVQTYNQLFEENPINASGVDA
KAILSARLSKSRRLENLIAQLPGEKKNGLFGNLIALSLGLTPNFKSNFDLAE
DAKLQLSKDTYDDDLDNLLAQIGDQYADLFLAAKNLSDAILLSDILRVNTEITKA
PLSASMIKRYDEHHQDLTLLKALVRQQLPEKYKEIFFDQSKNGYAGYIDGG
ASQEEFYKFIKPILEKMDGTEELLVKLNREDLLRKQRTFDNGSIPHQIHLGE
LHAILRRQEDFYPFLKDNREKIEKILTFRIPYYVGPLARGNSRFAWMTRKSEETI
TPWNFEEVVDKGASAQSFIERMTNFDKNLPNEKVLPKHSLLYEYFTVYNELT
KVKYVTEGMRKPAFLSGEQKKAIVDLLFKTNRKVTVKQLKEDYFKKIECFD
SVEISGVEDRFNASLGTYHDLLKIIKDKDFLDNEENEDILEDIVLTLTLFEDREM
IEERLKTYAHLFDDKVMKQLKRRRYTGWGRLSRKLINGIRDKQSGKTILDFL
KSDGFANRNFMQLIHDDSLTFKEDIQKAQVSGQGDSLHEHIANLAGSPAIKK
GILQTVKVVDELVKVMGRHKPENIVIEMARENQTTQKGQKNSRERMKRIEEGIK
ELGSQILKEHPVENTQLQNEKLYLYYLQNGRDMYVDQELDINRLSDYDVDAI
VPQSFLKDDSIDNKVLTRSDKNRGKSDNVPSEEVVKKMKNYWRQLLNAKLIT
QRKFDNLTKAERGGLSELDKAGFIKRQLVETRQITKHVAQILDSRMNTKYDEND
KLIREVKVITLKSKLVSDFRKDFQFYKVREINNYHHAHDAYLNAVVGTALIK
KYPKLESEFVYGDYKVYDVRKMIAKSEQEIGKATAKYFFYSNIMNFFKTEI
TLANGEIRKRPLIETNGETGEIVWDKGRDFATVRKVLSMPQVNIVKKTEVQTGGF
SKESILPKRNSDKLIARKKDWDPKKYGGFDSPTVAYSVLVVAKVEKGKSKK
LKSVKELLGITIMERSSFEKNPIDFLEAKGYKEVKKDLIIKLPKYSLFEL
ENGRKRMLASAGELQKGNELALPSKYVNFLYLASHYEKLKGSPEDNEQKQLFVEQ
HKHYLDEIIEQISEFSKRVILADANLDKVLSAYNKHRDKPIREQAENIIHL
FTLTNLGAPAAFKYFDTTIDRKQYRSTKEVLDATLIHQSITGLYETRIDLS
QLGGD
>sp|RETBP_HUMAN|
MKWVWALLLLAALGSGRAERDCRVSSFRVKENFDKARFSGTWYAMAKKDP
EGLFLQDNIVAEFSVDETGQMSATAKGRVRLLNNWDVCADMVGTFTDTED
PAKFKMKYWGVASFLQKGNDDHWIVDTDYDTYAVQYSCRLLNLDGTCADS
YSFVFSRDPNGLPPEAQKIVRQRQEELCLARQYRLIVHNGYCDGRSERNL
L
>sp|CAH1_HUMAN|
ASPDWGYDDKNGPEQWSKLYPIANGNNQSPVDIKTSETKHDTSLKPISVS
YNPATAKEIINVGHSFHVNFEDNDNRSVLKGGPFSDSYRLFQFHFHWGST
NEHGSEHTVDGVKYSAELHVAHWNSAKYSSLAEAASKADGLAVIGVLMKV
GEANPKLQKVLDALQAIKTKGKRAPFTNFDPSTLLPSSLDFWTYPGSLTH
PPLYESVTWIICKESISVSSEQLAQFRSLLSNVEGDNAVPMQHNNRPTQP
LKGRTVRASF
>sp|MYG_HUMAN|
GLSDGEWQLVLNVWGKVEADIPGHGQEVLIRLFKGHPETLEKFDKFKHLK
SEDEMKASEDLKKHGATVLTALGGILKKKGHHEAEIKPLAQSHATKHKIP
VKYLEFISECIIQVLQSKHPGDFGADAQGAMNKALELFRKDMASNYKELG
FQG
>sp|K1C15_SHEEP|
MATTLLQTSSSTFGGSSTRGGSLLAGGGGFGGGSLYGGGGSRTISASSAR
FVSSGSAGGYGGGFGGGAGSGYGGGFGGGFGGGFGSGFGDFGGGDGGLLS
GNEKITMQNLNDRLASYLEKVRALEEANADLEVKIRDWYQRQSPTSPERD

YSPYFKTTDELRDKILAAAIDNSRVILEIDNARLAADDFRLKYENEMALR
QSVEADINGLRRVLDELTLTKTDLEMQIESLNEELAYLKKNHEEEMKEFS
NQLAGQVNVEMDAAPGVDLTRVLSEMREQYEAMAEKNRRDAEAWFFSKTE
ELNKEVASNTEMIQTSKSEITDLRRTIQGLEIELQSQLSMKAGLESTLAE
TDGRYAAQLQQIQGLISSIEAQLSELRSEMEAQNQEYKMLLDIKTRLEQE
IATYHSLLEGQDARMAGIGTGEASLGGGGGGKVRINVEESVDGKVVSSRK
REI
>sp|CTRB_BOVIN|
CGVPAIQPVLSGLARIVNGEDAVPGSWPWQVSLQDSTGFHFCGGSLISED
WVVTAAHCGVTTSDVVVAGEFDQGLETEDTQVLKIGKVFKNPKFSILTVR
NDITLLKLATPAQFSETVSAVCLPSADEDFPAGMLCATTGWGKTKYNALK
TPDKLQQATLPIVSNTDCRKYWGSRVTDVMICAGASGVSSCMGDSGGPLV
CQKNGAWTLAGIVSWGSSTCSTSTPAVYARVTALMPWVQETLAAN
>sp|AMYS_HUMAN|
MKLFWLLFTIGFCWAQYSSNTQQGRTSIVHLFEWRWVDIALECERYLAPK
GFGGVQVSPPNENVAIHNPFRPWWERYQPVSYKLCTRSGNEDEFRNMVTR
CNNVGVRIYVDAVINHMCGNAVSAGTSSTCGSYFNPGSRDFPAVPYSGWD
FNDGKCKTGSGDIENYNDATQVRDCRLSGLLDLALGKDYVRSKIAEYMNH
LIDIGVAGFRIDASKHMWPGDIKAILDKLHNLNSNWFPEGSKPFIYQEVI
DLGGEPIKSSDYFGNGRVTEFKYGAKLGTVIRKWNGEKMSYLKNWGEGWG
FMPSDRALVFVDNHDNQRGHGAGGASILTFWDARLYKMAVGFMLAHPYGF
TRVMSSYRWPRYFENGKDVNDWVGPPNDNGVTKEVTINPDTTCGNDWVCE
HRWRQIRNMVNFRNVVDGQPFTNWYDNGSNQVAFGRGNRGFIVFNNDDWT
FSLTLQTGLPAGTYCDVISGDKINGNCTGIKIYVSDDGKAHFSISNSAED
PFIAIHAESKL
>sp|OVAL_CHICK|
MGSIGAASMEFCFDVFKELKVHHANENIFYCPIAIMSALAMVYLGAKDST
RTQINKVVRFDKLPGFGDSIEAQCGTSVNVHSSLRDILNQITKPNDVYSF
SLASRLYAEERYPILPEYLQCVKELYRGGLEPINFQTAADQARELINSWV
ESQTNGIIRNVLQPSSVDSQTAMVLVNAIVFKGLWEKAFKDEDTQAMPFR
VTEQESKPVQMMYQIGLFRVASMASEKMKILELPFASGTMSMLVLLPDEV
SGLEQLESIINFEKLTEWTSSNVMEERKIKVYLPRMKMEEKYNLTSVLMA
MGITDVFSSSANLSGISSAESLKISQAVHAAHAEINEAGREVVGSAEAGV
DAASVSEEFRADHPFLFCIKHIATNAVLFFGRCVSP
>sp|SRPP_HEVBR|
MAEEVEEERLKYLDFVRAAGVYAVDSFSTLYLYAKDISGPLKPGVDTIEN
VVKTVVTPVYYIPLEAVKFVDKTVDVSVTSLDGVVPPVIKQVSAQTYSVA
QDAPRIVLDVASSVFNTGVQEGAKALYANLEPKAEQYAVITWRALNKLPL
VPQVANVVVPTAVYFSEKYNDVVRGTTEQGYRVSSYLPLLPTEKITKVFG
DEAS
>sp|CRP_HUMAN|
MEKLLCFLVLTSLSHAFGQTDMSRKAFVFPKESDTSYVSLKAPLTKPLKA
FTVCLHFYTELSSTRGYSIFSYATKRQDNEILIFWSKDIGYSFTVGGSEI
LFEVPEVTVAPVHICTSWESASGIVEFWVDGKPRVRKSLKKGYTVGAEAS
IILGQEQDSFGGNFEGSQSLVGDIGNVNMWDFVLSPDEINTIYLGGPFSP
NVLNWRALKYEVQGEVFTKPQLWP
>sp|KRA3_SHEEP|
TGSCCGPTFSSLSCGGGCLQPRYYRDPCCCRPVSCQTVSRPVTFVPRCTR
PICEPCRRPVCCDPCSLQEGCCRPITCCPTSCQAVVCRPCCWATTCCQPV
SVQCPCCRPTSCQPAPCSRTTCRTFRTSPCC
